# Data-driven modelling captures dynamics of the circadian clock of *Neurospora crassa*

**DOI:** 10.1101/2022.01.24.477555

**Authors:** Amit Singh, Congxin Li, Axel C. R. Diernfellner, Thomas Höfer, Michael Brunner

**Affiliations:** Heidelberg University Biochemistry Center, Im Neuenheimer Feld 328, 69120 Heidelberg, Germany; Theoretical Systems Biology (B086) Deutsches Krebsforschungszentrum, Im Neuenheimer Feld 267, 69120 Heidelberg, Germany; Institute for Biomechanics, ETH Zurich, Leopold-Ruzicka-Weg 4, 8093 Zurich, Switzerland; Tissue and Tumor Microenvironments Group, Kennedy Institute of Rheumatology, University of Oxford, Oxford OX3 7FY, UK

## Abstract

Eukaryotic circadian clocks are based on self-sustaining, cell-autonomous oscillatory feedback loops that can synchronize with the environment via recurrent stimuli (zeitgebers) such as light. The components of biological clocks and their network interactions are becoming increasingly known, calling for a quantitative understanding of their role for clock function. However, the development of data-driven mathematical clock models has remained limited by the lack of sufficiently accurate data. Here we present a comprehensive model of the circadian clock of *Neurospora crassa* that describe free-running oscillations in constant darkness and entrainment in light-dark cycles. To parameterize the model, we measured high-resolution time courses of luciferase reporters of morning and evening specific clock genes in WT and a mutant strain. Fitting the model to such comprehensive data allowed estimating parameters governing circadian phase, period length and amplitude, and the response of genes to light cues. Our model suggests that functional maturation of the core clock protein Frequency (FRQ) causes a delay in negative feedback that is critical for generating circadian rhythms.

## Introduction

Circadian clocks orchestrate daily cycles of biochemical, physiological and behavioral processes. Anticipation and adaptation to recurring environmental changes is thought to improve the fitness of organisms (Dunlap, 1999; Ouyang et al., 1998; Young and Kay, 2001). Circadian clocks share specific characteristics that are crucial for their function: (1) Circadian clocks generate in the absence of external stimuli a self-sustaining rhythm of about 24 h. (2) They respond to recurring external stimuli such as changes in temperature, light, and nutrient sources to synchronize with the 24-hour environmental day-night cycle. (3) Circadian clocks are temperature compensated, such that the period length of circadian rhythms is over a broad range not significantly affected by the average daily temperature (Aschoff, 1965; Pittendrigh, 1954; 1960). Misalignment of the circadian clock and the environment contributes to several biochemical and physiological disorders including insomnia, mood disorder, diabetes, and cancer (Bass and Lazar, 2016; Bechtold et al., 2010; Challet, 2007; Hellweger, 2010).

Circadian clocks of eukaryotes are based on cellular transcriptional-translational feedback loops (TTFLs) regulating the expression of core clock genes as well as clock-controlled genes (Dunlap, 1999; Lee et al., 2000; Lowrey and Takahashi, 2000). In *Neurospora crassa*, the hetero-dimeric transcription activator White Collar Complex (WCC) and its inhibitor FFC, a complex containing Frequency (FRQ), FRQ-interacting RNA-helicase (FRH) (Cha et al., 2011; Guo et al., 2010) and casein kinase 1a (CK1a) (Gorl et al., 2001), are the core components of the TTFL (see below).

Previously, several mathematical models of the *Neurospora* circadian clock have been built on the basis of the core negative feedback loop constituted by the WCC and FRQ (FFC). Due to the unavailability of comprehensive experimental data these models uncovered and described principle properties of the clock in a rather generic manner and did, in most cases, not include detailed molecular interactions and mechanisms (Akman et al., 2008; Bellman et al., 2018; Dovzhenok et al., 2015; Francois, 2005; Gin et al., 2013; Hong et al., 2008; Leloup et al., 1999; Liu et al., 2019; Ruoff et al., 1996; Ruoff and Rensing, 1996; Tseng et al., 2012; Upadhyay et al., 2019, Upadhyay et al., 2020, Burt et al., 2021). In this study, we analyzed a *WT* and a mutant strain, *Δvvd* (Malzahn et al., 2010), which is compromised in its capacity to adapt to light. We collected a comprehensive set of clock-related data by measuring *in vivo* in constant darkness and in light-dark cycles the expression of luciferase reporters of the core clock gene *frq* and of *vvd*. In addition, we measured luciferase reporters of *conidial separation-1* (*csp-1*), which encodes a morning-specific clock-controlled repressor and one of CSP-1’s target genes, *fatty acid metabolim-3* (*fam-3*) (Sancar et al., 2011). The data warranted building a complex mathematical model with rather detailed molecular interactions. The data-driven model allowed us to estimate not only the expression phase but also the amplitude of rhythmically expressed genes and to uncover promoter-specific properties that determine their function in dark and light. Our approach demonstrates how high-resolution data sets can be used to more optimally exploit the theoretical capabilities of mathematical modeling.

## Results and Discussion

### Interaction network of the *Neurospora* clock

To understand how the manifold molecular interactions implicated in the circadian clock of *Neurospora crassa* control autonomous circadian oscillations and entrainment, we established an interaction network based on the available data (Figure 1): White Collar-1 (WC-1) and White Collar-2 (WC-2) are PAS (PER-ARNT-SIM) containing GATA-type zinc finger proteins, which constitute the heterodimeric White Collar Complex (WCC), the core transcription activator of the circadian clock of *Neurospora* (Crosthwaite et al., 1997; He et al., 2002; Linden and Macino, 1997; Ponting and Aravind, 1997; Talora et al., 1999). In the dark, transcription of *wc-1* and *wc-2* are controlled by unknown TFs (Kaldi et al., 2006). We modeled transcription of *wcc* being equivalent to the transcriptional production of its limiting component *wc-1*. The subsequent translation and assembly of the WC-1 and WC-2 subunits constituting the WCC was described by a single production term.

**Fig. 1.**
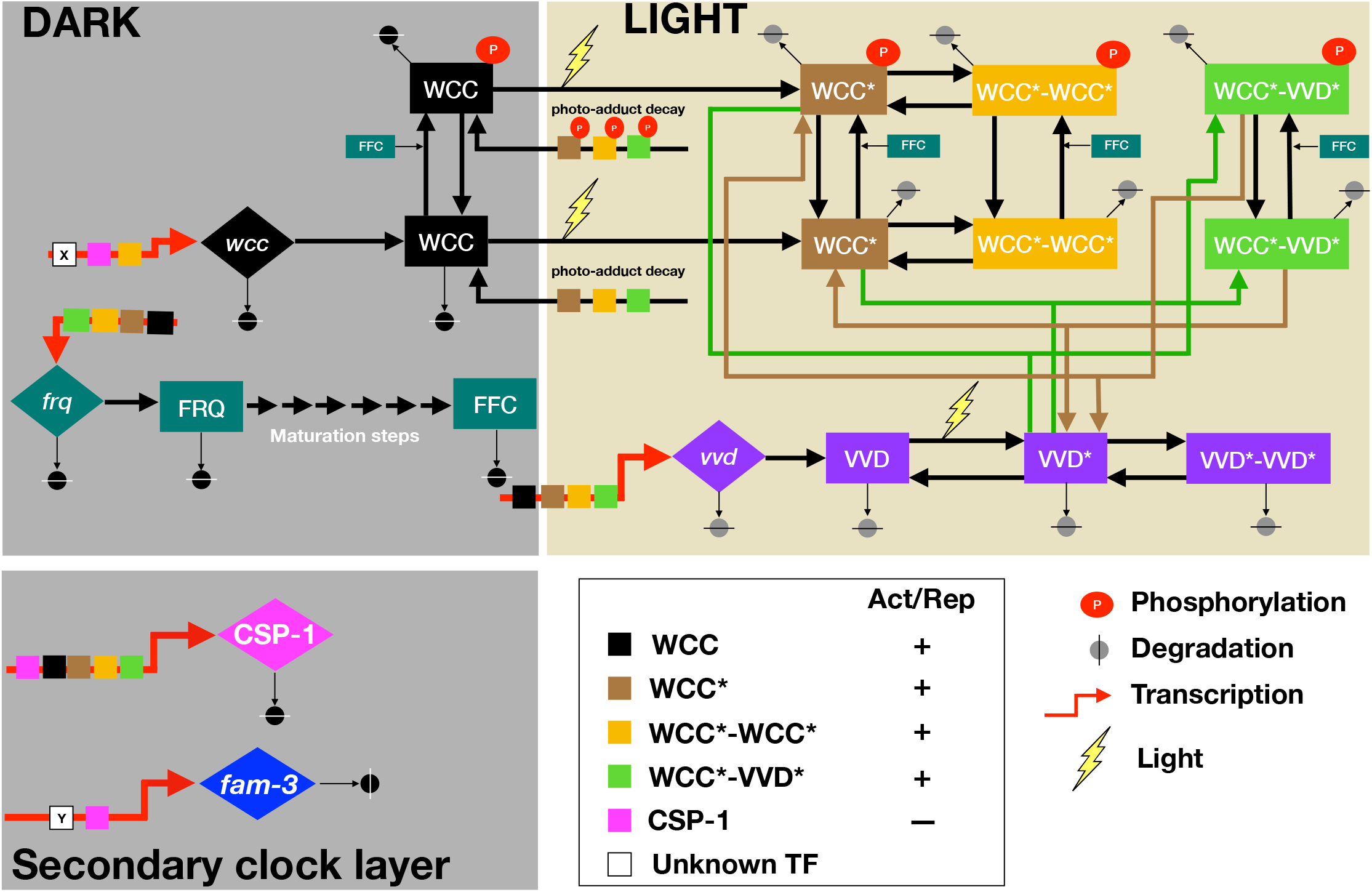
Schematic of the *Neurospora* circadian clock in dark and light. Thick red arrows indicate transcription, and the small colored square boxes on these arrows indicate the TFs controlling the respective gene. White boxes X and Y indicate unidentified TFs activating transcription of *wcc* and *csp-1*, respectively. Large diamond and square boxes represent the indicated mRNA and protein species, respectively. FRQ, inactive Frequency protein; FFC, assembled, active FRQ-FRH-CK1a complex; WCC, White Collar Complex, VVD, Vivid; WCC* and VVD*, light-activated species; *csp-1, conidial separation 1; fam-3, fatty acid desaturase*.

The WCC controls transcription of the core clock gene *frequency* (*frq*) (Aronson et al., 1994). FRQ protein was modeled as being initially inactive and unstable. FRQ homo-dimerizes (Cheng et al., 2001a), assembles with FRH (Cheng et al., 2005), and CK1a (Gorl *et al*., 2001), forming the active FFC complex (Shi et al., 2010). The FFC and/or individual subunits or subcomplexes shuttle into the nucleus, receiving potentially licensing phosphorylation by CK1a (Gorl *et al*., 2001; Querfurth et al., 2007) and/or by other kinases (Diernfellner and Brunner, 2020; Huang et al., 2007; Mehra et al., 2009; Yang et al., 2003; Yang et al., 2002; Yang et al., 2001). As little molecular detail is known on assembly and maturation, we modeled these processes in a generic manner by a linear chain of six consecutive steps. The FFC is the core negative element of the TTFL. It interacts transiently with and inactivates the WCC by facilitating its phosphorylation by the CK1a subunit of the FFC (Schafmeier et al., 2005), and hence, we modeled the FFC acting enzymatically on the WCC and converting it into its phosphorylated, inactive form, P-WCC. As the WCC controls morning-specific transcription of *frq*, its inactivation closes the circadian negative feedback loop.

Inactivated phosphorylated WCC is stable and accumulates at elevated levels, and, hence, FRQ is part of a positive feedback loop with respect to WCC abundance (Cheng et al., 2001b; Crosthwaite *et al*., 1997; Denault et al., 2001; Dunlap, 1999; Lee *et al*., 2000; Neiss et al., 2008; Shi *et al*., 2010).

In the course of a circadian period FRQ is progressively phosphorylated triggering its inactivation and degradation (Gorl *et al*., 2001; Larrondo et al., 2015). Inactivation of the FFC and degradation of its FRQ subunit was described by a single degradation/irreversible inactivation rate of the FFC. With decreasing amounts of active FFC, WCC is reactivated by dephosphorylation (Cha et al., 2008; Schafmeier *et al*., 2005) and replenished by de novo synthesis, and then a new circadian cycle begins with reactivation of *frq* transcription.

Our wiring schematic of the core circadian oscillator in the dark (Figure 1, upper left box) is topologically equivalent to the Goldbeter limit cycle oscillator model (Goldbeter, 1995). However, the Goldbeter model as well as other generic circadian oscillator models (Hong *et al*., 2008; Leloup *et al*., 1999; Ruoff *et al*., 1996; Ruoff and Rensing, 1996; Upadhyay *et al*., 2019) used a Hill-function for the transcriptional production of the core circadian inhibitors, PER in animals and FRQ in fungi, respectively, with Hill coefficients ranging from 2 to 7 (Bellman *et al*., 2018; Hong *et al*., 2008; Liu *et al*., 2019; Tseng *et al*., 2012). While the Hill-coefficients in these models helped generate robust oscillations, molecular interactions underlying such highly cooperative processes are not known. To attain smaller Hill exponents several previous models introduced a time delay by including intermediate steps in the negative feedback loop (Arcak and Sontag, 2006; Forger, 2011; Thron, 1991). A natural time delay is provided by the maturation of newly synthesized, inactive FRQ to active FFC, allowing us to describe WCC-activated transcription of *frq* by a simple Michaelis-Menten-like equation without introducing a Hill-coefficient.

The WCC controls rhythmic expression of many clock-controlled genes, among them *vivid*(*vvd*) and *conidial separation-1* (*csp-1*) (Froehlich et al., 2002; Malzahn *et al*., 2010; Sancar et al., 2015; Smith et al., 2010), which were included in our model.

Light cues directly activate the WCC and thereby reset the circadian clock and induce several cellular processes including biosynthesis of carotenoids, asexual conidiospores formation, and development of sexual structures (Corrochano, 2007; Harding and Turner, 1981; Lauter and Russo, 1991; Ruger-Herreros et al., 2014; Sancar *et al*., 2015; Sancar *et al*., 2011). The light-activated WCC is a potent transcription activator of *frq and wc-1* (*wcc* in our model) as well as of many light-responsive genes including *vvd* and *csp-1* (Chen et al., 2009; Froehlich *et al*., 2002; Malzahn *et al*., 2010; Sancar *et al*., 2015).

The WC-1 subunit of the WCC and VVD are blue-light photoreceptors harboring a flavin-binding light-oxygen-voltage- (LOV) domain (for review see (Demarsy and Fankhauser, 2009; Suetsugu and Wada, 2013)). VVD has no known function in the dark. Upon exposure to blue-light, a photo-adduct between a conserved cysteine residue of the LOV-domain and its bound FAD cofactor is formed (Zoltowski and Crane, 2008). WC-1 is activated in corresponding fashion (Cheng et al., 2003; Malzahn *et al*., 2010). The photo-adducts stabilize a conformation of the LOV domains that favors highly dynamic homo- and heterodimerization of VVD* and WCC* (Malzahn *et al*., 2010; Zoltowski and Crane, 2008). The light-activated WCC homodimer (WCC*WCC*) binds to light-response elements (LREs) (Froehlich *et al*., 2002). WCC*WCC* was modeled as a potent transcription activator of *frq, vvd, wcc* (*wc-1*), and *csp-1* (Chen *et al*., 2009; Froehlich *et al*., 2002; Malzahn *et al*., 2010; Sancar *et al*., 2015; Smith *et al*., 2010). As the interaction of VVD* with WCC* leads to photoadaptation of light-dependent transcription, the activity of the WCC*VVD* heterodimer and of monomeric light-activated WCC* was modeled as being equivalent to the activity of the dark form of the WCC. Light-activation triggers rapid hyperphosphorylation and accelerated degradation of WCC* (Kaldi *et al*., 2006; Linden and Macino, 1997; Schafmeier et al., 2008; Schafmeier and Diernfellner, 2011) and thereby also affects levels of its readily equilibrating complexes, WCC*WCC* and WCC*VVD*. The light-activated, unphosphorylated WCC* is unstable and rapidly degraded (Kaldi *et al*., 2006; Linden and Macino, 1997; Schafmeier *et al*., 2008; Schafmeier and Diernfellner, 2011), while interaction with VVD* stabilize the WCC*. The light-induced phosphorylation of WCC* was not explicitly modeled but is included into the higher degradation rate of light activated species, which was constraint in our model to a half-time ≤ 3 h.

In contrast, the FRQ dependent phosphorylation, inactivation and stabilization of all WCC species (Schafmeier *et al*., 2005) was explicitly modeled. The LOV-domain photo-adducts of WCC* and VVD* decay spontaneously into their dark forms with a half-time in the range of hours (Malzahn *et al*., 2010; Zoltowski and Crane, 2008).

All molecular reactions in the model were translated to a mathematical form describing either Michaelis-Menten or simple kinetics equations. The model contains 18 variables including mRNA and protein species. The detailed mathematical equations are displayed in Supplemental Information. The model includes the key features of the circadian clock and is sufficiently complex to allow an adequate, and mostly quantitative, molecular interpretation of experimental data.

### High-amplitude response to light versus low-amplitude dark oscillations

In order to collect sufficient data as a basis for the molecular model of the circadian clock we generated *WT* and *Δvvd* reporter strains expressing destabilized luciferase (lucPEST) (Cesbron et al., 2013) under the control of the morning-specific *frq, vvd* and *csp-1* promoters and the evening-specific *fam-3* promoter. The *lucPEST* transcription units carried the 3’ region of the *trpC* gene of *Aspergillus nidulans* for termination of transcription and the reporter genes were inserted downstream of the *his-3* locus into the genome of *Neuropora*. Mycelial cultures of these strains were grown in 96-well plates (with more than 30 replicates in 3 independent experiments) on solid agar medium. The medium contained sorbose in order to restrict growth, which allows live recordings of bioluminescence over many days. The cultures were grown on the sorbose medium and synchronized by exposure to 12 h light, 12 h dark and 12 h light and then transferred to dark for 24 h before the luciferase measurement was started (t=0) (Figure 2). After 4 days in constant darkness (t = 72 h) the samples were exposed to 12 h light, 12 h dark and 12 h light, and then kept in the dark for another 64 h. In constant darkness the expression levels of all *lucPEST* reporters oscillated but showed dampening over time (see Figure 2). It is not clear whether and to what extent the dampening is the consequence of desynchronization of individual nonconnected hyphae or due to a real reduction of amplitude of the oscillator.

**Fig. 2.**
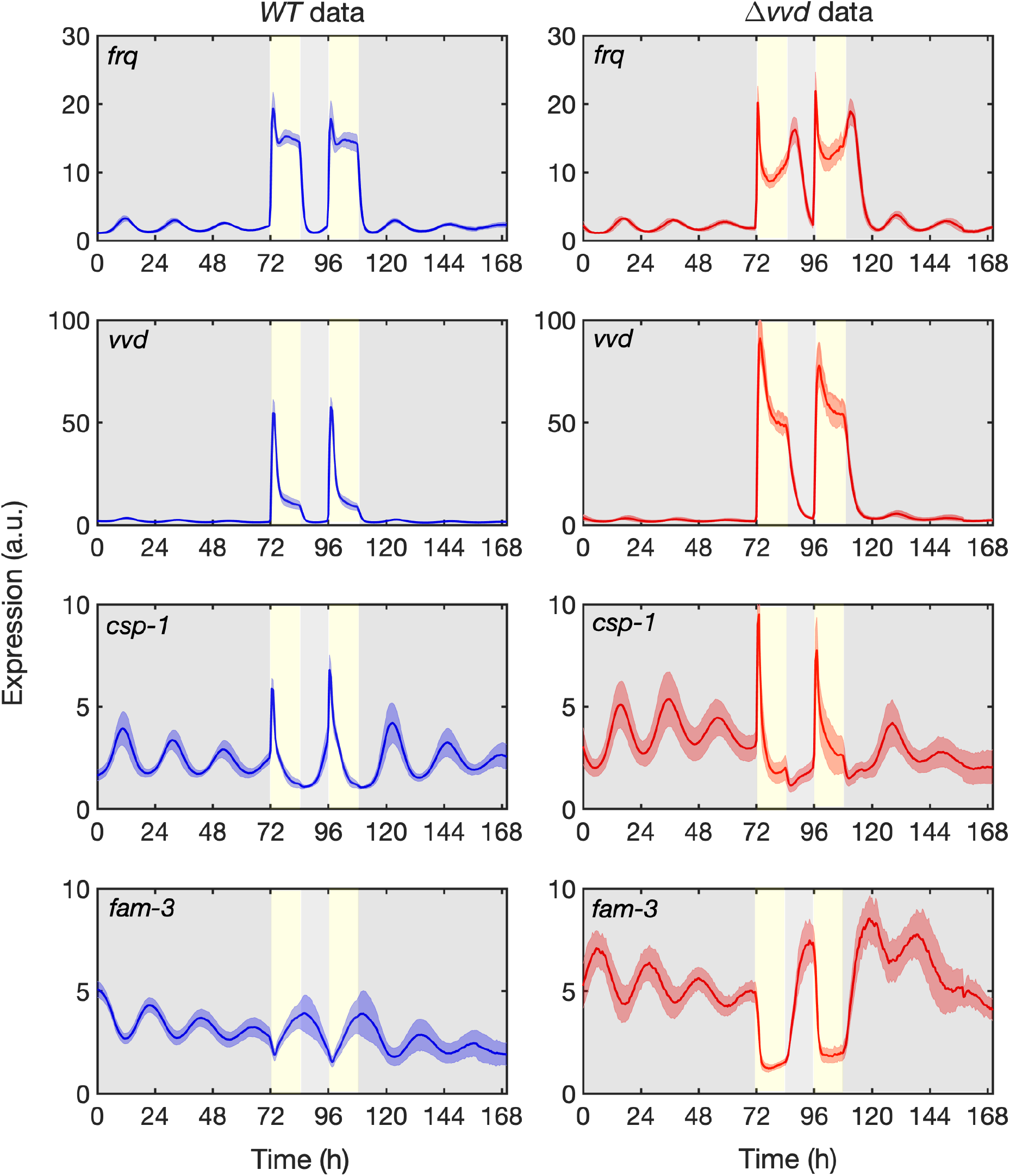
Temporal expression profiles of reporter genes in DD and LD. Luciferase activity under the control of the indicated promoters in *WT* (left panels) and *Δvvd* (right panels). The solid line represents the average of 30 measurements from 3 independent experiments. The shaded areas correspond to the standard deviation, SD. Yellow vertical boxes and grey areas indicate 12 h light periods and dark periods, respectively.

The expression of *frq, vvd*, and *csp-1* reporters oscillated with peaks levels in the subjective morning, while *fam-3*, which is repressed by CSP-1 in the morning, was rhythmically expressed in antiphase with peak levels in the subjective evening. As expected on the basis of previous studies (Cesbron *et al*., 2013; Elvin et al., 2005; Heintzen et al., 2001; Hunt et al., 2007), all oscillations were phase-delayed in *Δvvd* (Figure 2, right panels). After the light was turned on, expression of *frq* was rapidly induced in *WT* to a level ~18-fold higher, but then dropped and adapted to a level ~12-fold higher than in the dark. Expression of *vvd* was rapidly induced ~50-fold compared with the level in the dark and then adapted to a ~10-fold higher level, and csp-1 was induced ~6-fold and fully adapted to its dark expression level. Thus responses of *frq, vvd*, and *csp-1* to light had much larger amplitudes than autonomous oscillations in constant darkness. Moreover, the three light-inducible promoters responded with different inactivation kinetics to light-activated WCC.

In contrast, the evening-specific *fam-3* reporter was transiently repressed after the light was turned on. This is consistent with the light-induced transient expression of CSP-1, which represses its own gene, *csp-1*, as well as *fam-3* and many other genes (Sancar *et al*., 2011).

In *Δvvd*, the circadian oscillation of all reporters was phase-delayed (arrows in Figure 2) as expected. The transcriptional dynamics in light-dark cycles of *frq* and *vvd*were quite different in *Δvvd*as compared to *WT*. The light response of the *vvd* reporter was qualitatively similar to that in *WT*. However, the spike level after light on was almost twice that of *WT*, and the light-adapted expression level was ~5-fold higher than in *WT*. These data are consistent with the increased activity of light-activated WCC in the absence of its light-dependent inhibitor, VVD (Malzahn *et al*., 2010). In contrast, the initial light-induced spikes of the *frq* reporter were similar in *Δvvd* and *WT*, suggesting that the *frq* promoter was already functionally saturated at *WT* levels of light-activated WCC. After the initial light-induced spike, *frq-lucPEST* expression levels decreased sharply and more markedly in *Δvvd* than in *WT*, and then increased again in the further course of the light phase. When light was turned off, *vvd* transcription decreased as expected, consistent with the decreasing level of light-activated WCC. Surprisingly, however, the *frq* level in *Δvvd* increased transiently after the lights were turned off and then dropped. The difference in expression dynamics of *frq* compared with *vvd* after light on and even more so after the light-to-dark transition is consistent with the previously reported refractory behavior of the *frq* promoter (Cesbron et al., 2015; Li et al., 2018). Indeed, in *Δvvd* the *frq* promoter is partially repressed in light, and hence not maximally active. The rapid decrease of the light-induced transcription spike after light on reflects the dynamics of the light-dependent partial repression. Similarly, the transient increase in *frq-lucPEST* transcription in *Δvvd* after turning off the light is consistent with the repression of the *frq* promoter being relieved faster than the WCC activity (level) decreases.

### The model quantitatively captures circadian oscillations and light entrainment in *WT Neurospora*

Our model was trained to the temporal expression profiles of the *frq, vvd, csp-1* and *fam-3* reporters in *WT* and *Δvvd*. The parameter space was restricted to a biological meaningful range as described in Supplemental Information. The parameters were estimated using maximum likelihood (lsqnonlin optzimizer implemented in D2D/Matlab; (Raue et al., 2015; Sancar *et al*., 2011)).

Overall, our model simulations show oscillation of all reporters in the dark and responded in appropriate manner to the LD-cycles (Figure 3). The simulations of *frq* and *vvd* transcription in *WT* captured the experimental data quite well, reproducing period length, phase and amplitude of the free-running oscillations as well as the transcription dynamics in LD cycles (Figure 3, left panels). The simulations of *csp-1* expression in *WT* captured period length and phase in DD and part of the dynamics in LD cycles, while amplitude of the model deviated somewhat from the measured data. In our model, we assumed that *csp-1* is rhythmically activated by the WCC and rhythmically repressed by CSP-1. The deviation of data and model could indicate that, in addition to WCC, an unknown transcription activator contributes to the *csp-1* expression. Furthermore, compared to the luciferase data, the model predicted slightly higher *csp-1* levels towards the end of the 12 h light phases, and the level then dropped abruptly after lights-off in the model. This sudden drop in *csp-1* RNA, and hence of the short-lived CSP-1 repressor (Sancar *et al*., 2011), lead to a rapid relieve of *fam-3* repression and thus, the model showed transiently slightly higher *fam-3* levels than experimentally observed.

**Fig. 3.**
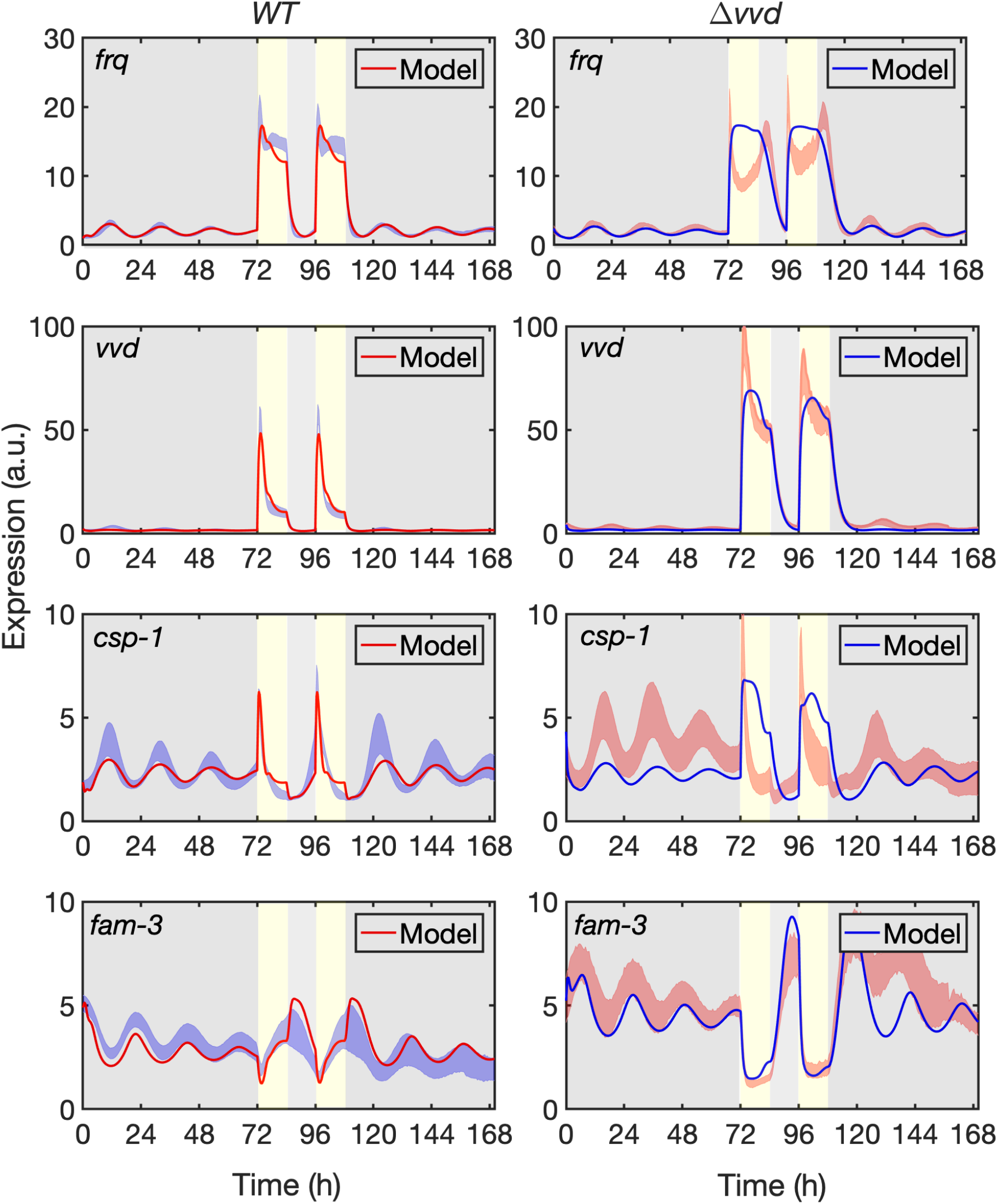
Model fitted to reporter gene expression in DD and LD. Trajectories of the model (solid red lines in *WT*and solid blue lines in *Δvvd*) fitted to the luciferase activity expressed under the control of the indicated promoters (standard deviation is shown, see Figure 2) in *WT* (left panels) and *Δvvd* (right panels). Yellow vertical boxes and grey areas indicate 12 h light periods and dark periods, respectively.

The simulations of gene expression in *Δvvd* (Figure 3, right panels) faithfully reproduced the period length and phase of all reporters and captured their phase delays compared with *WT* (Figure S1). The amplitudes of *frq* and *vvd* oscillations in the dark were modeled quite well. In contrast the amplitude of the *csp-1* oscillation was underestimated even more than in *WT*, supporting the notion (see above) that the *csp-1* transcription could be additionally supported by an unknown activator. The expression dynamics of *vvd* in LD cycles were fully captured, suggesting that Michaelis-Menten kinetics is suitable to quantitatively describe the activity of the *vvd*promoter in *WT*and in *Δvvd*. The higher light-induced expression level of the *vvd* reporter in *Δvvd* compared with *WT* was also captured by our model. Because the transcriptional output of the *vvd* promoter is very sensitive to the activity of the light-activated WCC (Li *et al*., 2018), the agreement of our model with the *Δvvd* and *WT* data justifies our differentiation of various light-activated WCC species (WCC*, WCC*-WCC* and WCC*VVD*) and the rates of their interconversion. Our model did, however, not capture the complexity of the expression profile of *frq* in light-dark cycles, indicating that Michaelis-Menten-like promoter activation was not sufficient to describe the response of the *frq* promoter to light cues, in particular in *Δvvd*.

Specifically, in *Δvvd*, our Michaelis-Menten-based model of the *frq* promoter produced almost square-waves of *frq* expression in LD with a rather slow decline after light was turned off. In contrast, the luciferase measurements revealed an initial overshoot of *frq* expression after light on, followed by rapid adaptation of *frq* in continuous light and then a transient increase in *frq* expression after the LD transition. As discussed above, the complex transcription dynamics reflect the previously reported partial repression of the light-activated *frq* promoter (Cesbron *et al*., 2015; Li *et al*., 2018). This particular feature of the *frq* promoter has not yet been included in our molecular model scheme as the underlying mechanism is still awaiting identification and functional characterization of the components involved. However, it is only the acquisition of high-resolution data that made it possible to reveal the discrepancy between model and data and thus predict that the transcriptional dynamics of the *frq* promoter in response to light cannot be modeled by simple Michaelis-Menten-like promoter activation. Such prediction is not possible with generic models.

Expression of *csp-1* in the *Δvvd* background increased sharply after light on and then adapted rapidly, reflecting the negative feedback of CSP-1 on its own transcription. The dynamics of the CSP-1 feedback, which was captured by the model in *WT*, was underestimated in *Δvvd*, indicating that the model did not quantitatively describe the enhanced light-induced expression of CSP-1 in absence of VVD. Potentially, Michaelis-Menten-like promoter regulation is not sufficient to describe the dynamics of the *csp-1* promoter.

We then challenged the model by simulating 10 consecutive light-dark cycles (Figure S2, left panels). The system responded in the same manner to each of the LD cycles. None of the components was depleted or accumulated to higher levels over the time period, indicating that the system was balanced.

In constant darkness the expression levels of all *lucPEST* reporters oscillated but showed dampening over time (see Figure 2). Our model was trained on the dampening bioluminescence oscillations in the dark it reproduced the dampening (Figure 3). Indeed, simulating prolonged incubation in the dark led to a substantial loss of amplitude (Figure S2, right panels). Since we have not allowed the possibility of desynchronization of individual oscillators in our analytical model, the damping in our model is due exclusively to a gradual loss of amplitude. However, it is possible that the experimentally observed dampening is due to desynchronization and that the modelled loss of amplitude does not reflect a physiological relevant process. Indeed, loss of amplitude was prevented when the dephosphorylation rate of WCC, i.e. its reactivation, was slightly increased (Figure S3).

Overall, however, the model almost quantitatively predicted the transcriptional dynamics of two hierarchical levels of the circadian clock in the dark and in light-dark cycles in *WT*. The model was generally less precise in predicting transcription rhythms and dynamics in *Δvvd*.

The detailed data-based model allowed us to ask specific questions that cannot be addressed with a generic clock model. For example, previous models of the *Neurospora* circadian clock used Hill functions which was crucial for robust circadian oscillation of *frq* transcription (Bellman *et al*., 2018; Dovzhenok *et al*., 2015; Leloup *et al*., 1999; Liu *et al*., 2019; Ruoff and Rensing, 1996). As we did not introduce a Hill-function for the transcriptional production of *frq*, the circadian oscillation in our model depends critically on the delay between the synthesis of *frq* and the appearance of the fully assembled and active FFC. This process, which was modeled by FRQ translation and six generic maturation steps, introduced a delay in the accumulation of active FFC and supported circadian oscillation of *frq*. Shortening the delay by successive removal of maturation steps resulted in increasing expression levels of *frq* and arrhythmicity (Figure 4A). Thus, our model suggests that maturation of the FFC may be an important process that should be studied experimentally. Indeed, this aspect is still very poorly understood, and we do not know how, when, in what order, and in what cellular compartments FRQ dimerizes, assembles with FRH and CK1a, and if FRQ requires licensing phosphorylation somewhere along this pathway to become an active inhibitor of WCC in the nucleus.

**Figure 4.**
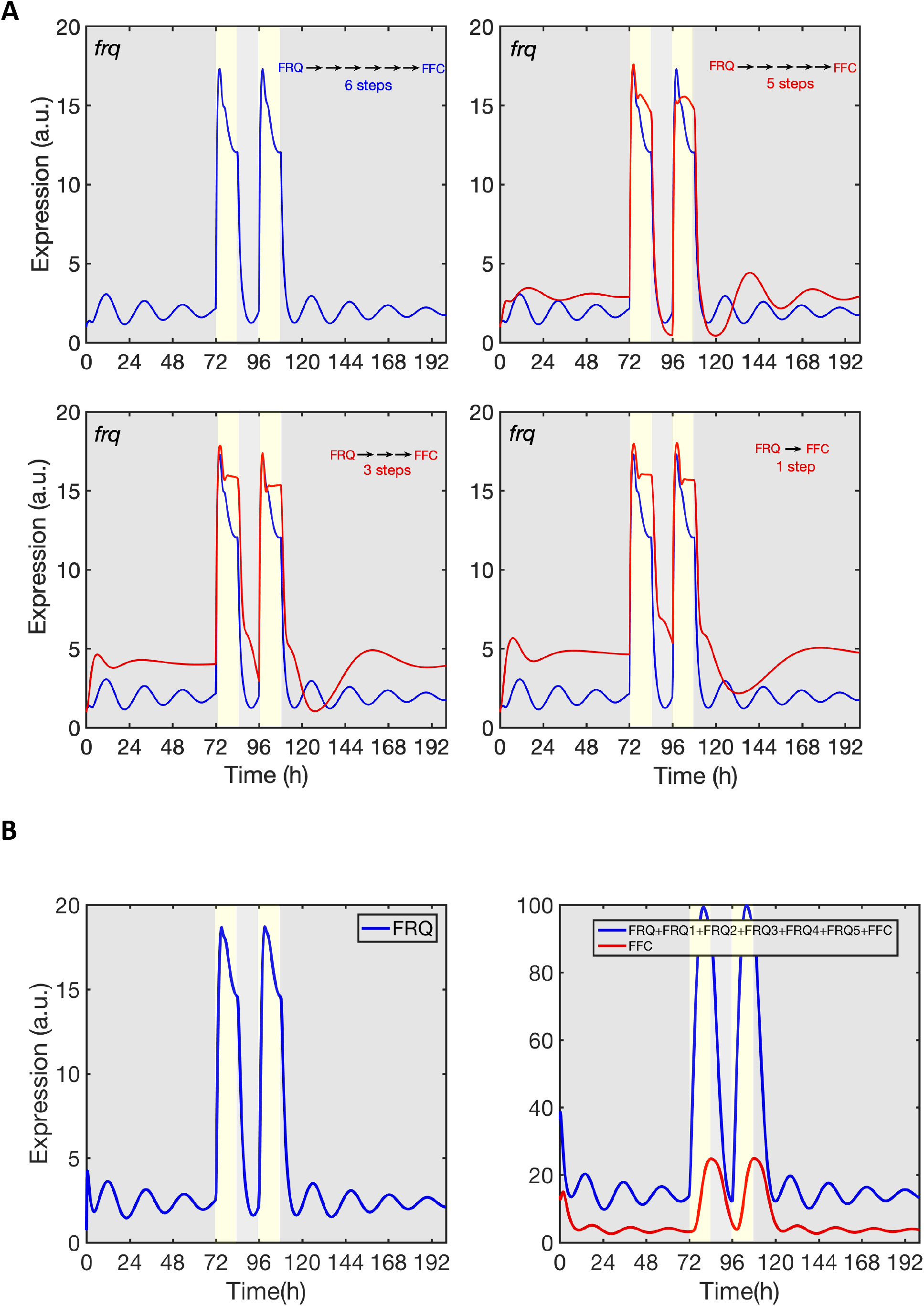
Maturation of inactive FRQ to active FFC is crucial for circadian rhythmicity. (A) Maturation of FRQ to FFC was modelled by a linear chain of 6 generic steps. The impact of the number of steps on *frq* expression levels and rhythm is shown. (B) Delay between newly synthesized FRQ and assembled FRH (left panel) and abundance of all unassembled FRQ-species versus assembled FRH (right panel).

Our model also predicts that only a fraction of the newly synthesized FRQ assembles with FRH while the majority is degraded. Due to the slow assembly process unassembled and partially assembled FRQ is more abundant than fully assembled, active FFC at any given time (Figure 4B). The high fraction of partially assembled FRQ species could explain the substoichiometic levels of FRH that are found in complex with total FRQ *in vivo* (Hurley et al., 2013), although one molecule of FRQ is capable of binding one molecule of FRH (Lauinger et., 2014).

The continuous recording of promoter-specific luciferase reporters *in vivo* in dark and light made it possible to estimate the intrinsic firing rate of the *frq* and *vvd* promoters (v_max_) and to uncover functional differences in promoter architecture. Previous quantitative ChIP-PCR analyses revealed a similar affinity of the light-activated WCC for the LREs of *frq* and *vvd* (Cesbron *et al*., 2015; Li *et al*., 2018). Yet, our model predicted a small K_M_ for WCC-dependent transcription activation of *frq* and a larger K_M_ for *vvd*. The data indicate that the affinity of the transcription factor for its LRE does not directly correlate with gene transcription. The difference likely reflects that the *frq* LRE is located in the core promoter such that bound WCC*WWC* can directly activate the core promoter. In contrast, the *vvd* LRE is located in an upstream enhancer region. Hence, activation of *vvd* transcription is dependent on looping of the LRE-bound WCC*WWC* to the core promoter. The coupled equilibria of TF-binding to the remote LRE and looping of the LRE-bound TF to the promoter result in an overall higher K_M_ for the activation of transcription at the *vvd*promoter (Li *et al*., 2018). Furthermore, our model predicted a small v_max_ for *frq*, consistent with *frq* being a weak promoter with an intrinsically low firing rate. The light-activated *vvd* promoter has a high intrinsic firing rate, consistent with a large predicted v_max_. Our simulations with these promoter-specific parameters (Figure 5A, B) captured in principle the previously reported saturation of *frq* transcription at rather low light intensity while *vvd* responds over a much wider range proportional to the intensity of light (Cesbron *et al*., 2015; Li *et al*., 2018).

**Figure 5.**
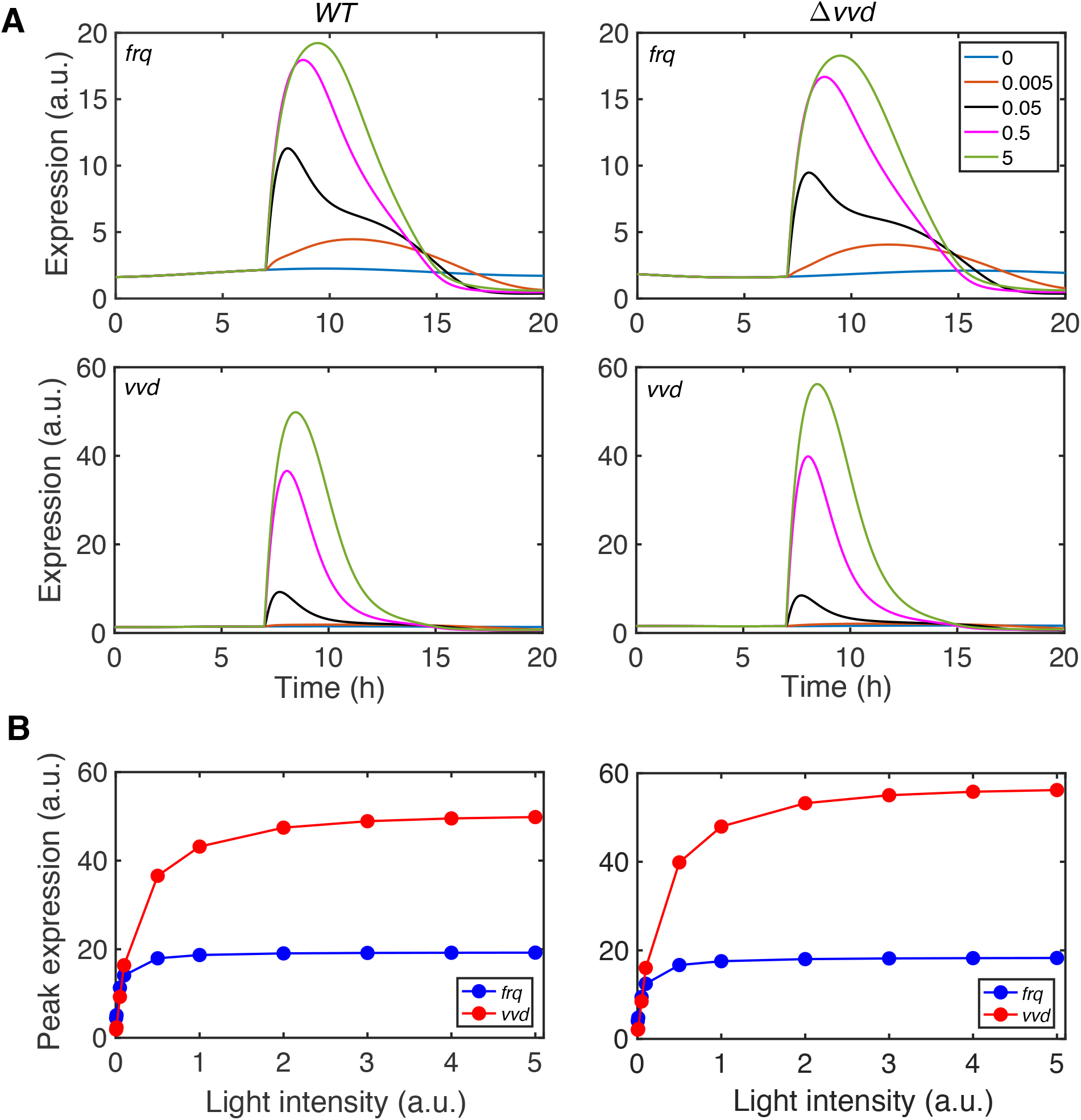
Modelling the response of *frq* and *vvd* promoters to light-pulses of different intensity. *WT* (left panels) and *Δvvd* (right panels) were exposed at t = 7 min to a virtual LP. (A) Modeled response of *frq* and *vvd* to virtual LPs of the indicated intensities. (B) Plot of peak expression levels of *frq* and *vvd* versus LP intensity. Simulated LP intensities: 0; 0.005; 0.005; 0.05; 0.5 and 5.0 arbitrary units (a.u.).

In summary, the *luc* reporter strains allowed measuring transcript levels and dynamics with high temporal resolution. Obtaining such comprehensive data justified the construction of a detailed molecular model of the *Neurospora* circadian system. The wiring scheme of the model and the derived kinetic parameters captured crucial features of the circadian system, but cannot yet accurately predict the temporal dynamics of all processes. This discrepancy led to new testable hypotheses. Overall, therefore, we believe that data-based approaches to circadian clocks as the one presented here can become a valuable tool for quantitatively understanding the interaction of the molecular clock components.

## Acknowledgements

The authors gratefully acknowledge the data storage service SDS@hd supported by the Ministry of Science, Research and the Arts Baden-Württemberg (MWK) and the German Research Foundation (DFG) through grant INST 35/1314-1 FUGG and INST 35/1503-1 FUGG. M.B. and T.H. are supported by the Deutsche Forschungsgemeinschaft, TRR186.

## Author Contributions

M.B. and T.H. conceived the study. A.S. and C.L. performed the modelling analysis. A.C.R.D. carried out the experiments. A.S., M.B. wrote the paper.

## Materials and methods

### *Neurospora* strains and plasmids

The *Neurospora* strains denoted with *WT* and *Δvvd* carried the *ras1^bd^* mutation (Belden et al., 2007) and either a *pfrq-lucPEST, pvvd-lucPEST* (Cesbron *et al*., 2013), *pfam3-lucPEST*, formerly called desat-lucPEST (Sancar *et al*., 2011) or a p*csp1-lucPEST* reporter gene integrated downstream of the *his-3* locus. For p*csp1-lucPEST*, a 7395bp fragment immediately upstream of the csp-1 ORF was amplified from gDNA and cloned in front of the lucPEST ORF using EcoRI (vector)/MfeI(PCR insert) and NotI restriction sites. The resulting plasmid was used to transform the above mentioned *Neurospora* strains.

### Real-time luciferase activity measurements

Sorbose medium containing 1x FGS (0.05 %fructose, 0.05% glucose, 2% sorbose), 1x Vogels, 1% agarose, 10 ng/ml biotin, and 25 μM firefly Luciferin was used for the assessment of the luciferase activity. 96-well plates were inoculated with 3 X 10^4^ conidia per well and incubated in DD at 25°C. Bioluminescence was recorded in DD or LD at 25° C with EnSpire Multilabel Readers (Perkin Elmer). The light intensity was 0.25 *μ*E. Three independent experiments with multiple biological replicates each were performed to generate the data (n≥30).

### Mathematical modelling and parameter estimation

The reactions of the model were translated to ODEs

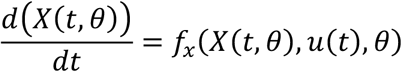

where ***θ*** is a parameter ***θ*** = (***θ*_1_, *θ*_2_.. *θ_i_***). The initial state of the system is described by ***X***(**0, *θ***) = ***f*_*X*_0__**(***θ***) The variables ***X*** correspond to the dynamics of the concentration of molecular components of the model. To derive the unknown model parameters, the circadian model was calibrated by a maximum likelihood estimation using quantitative experimental data obtained by luciferase measurements. The model was fitted to the luciferase data using (MATLAB version 2016b) D2D software package from http://www.data2dynamics.org (Raue et al. 2015).

---

**Fig. S1.**
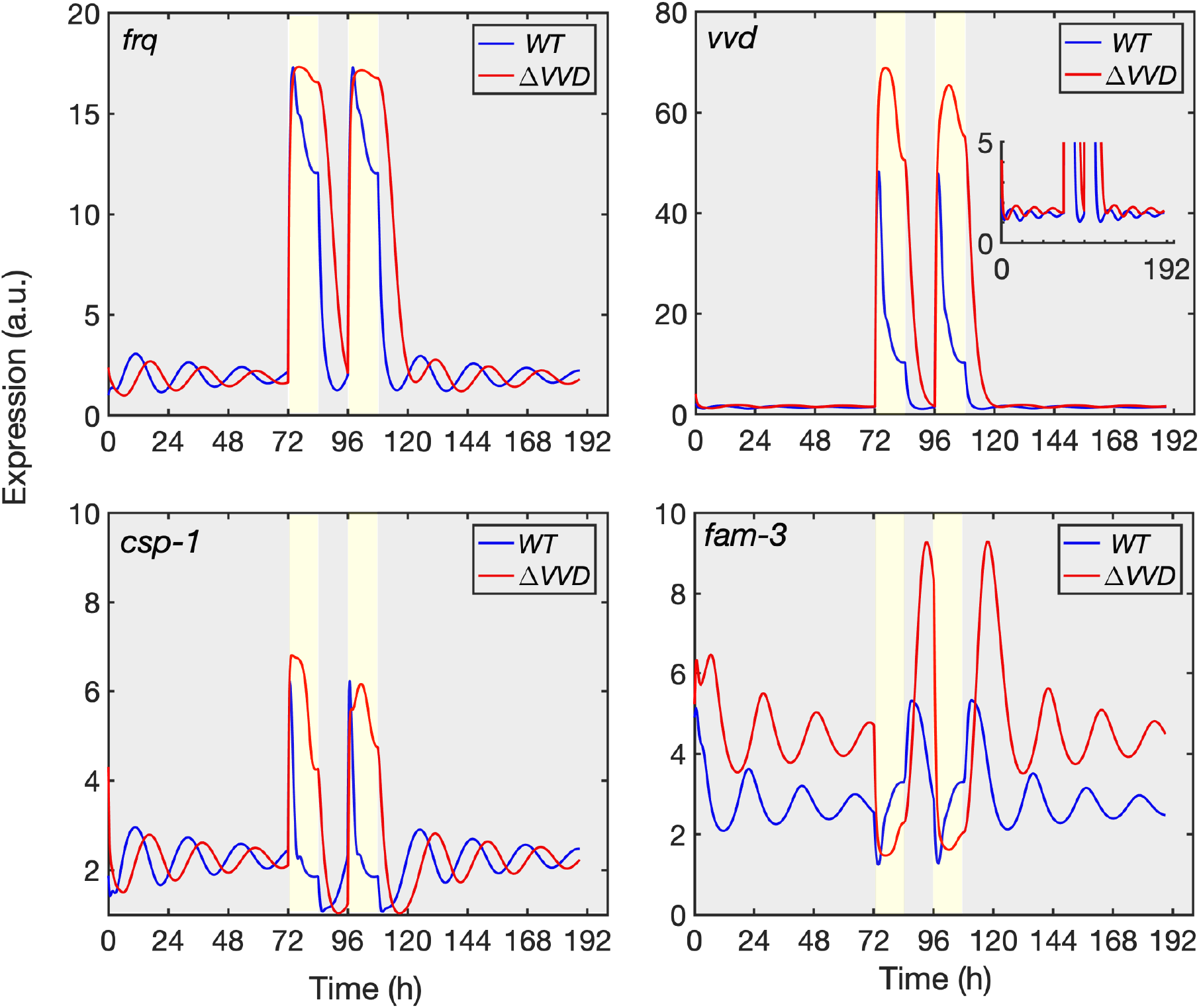
Phase delay of expression rhythms in *Δvvd*. Modelled trajectories of the expression rhythms in WT and *Δvvd* (see Figure 3) were superimposed.

**Fig. S2.**
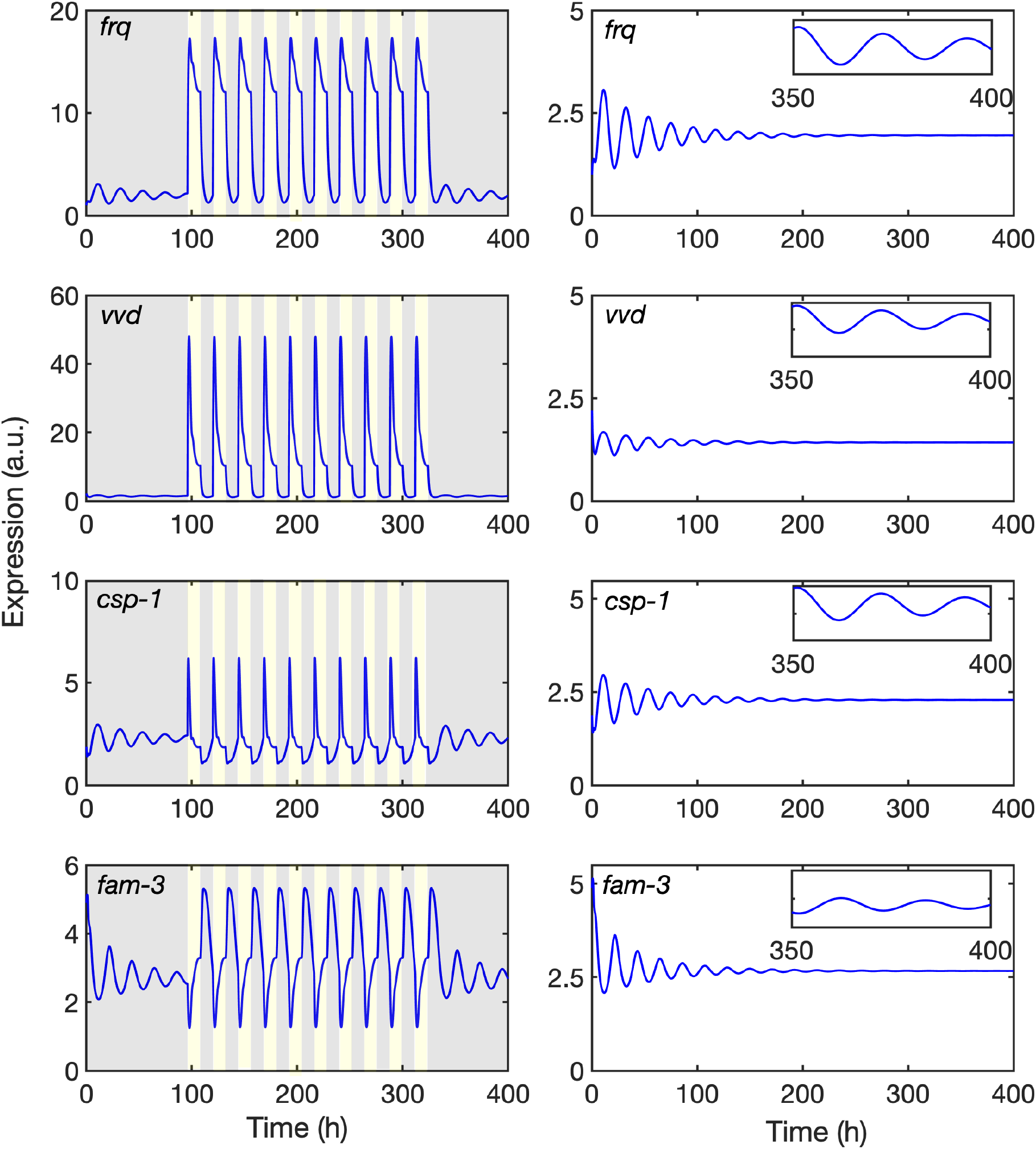
Characterization of the model. (**A**) Simulated response of reporters to 10 repetitive LD cycles. (**B**) Rhythms of reporter genes dampen in constant darkness. Expression rhythms in the dark of the reporters were modelled for 400 h. Inserts show zoom-in of the expression rhythms between 350 and 400 h.

**Fig. S3.**
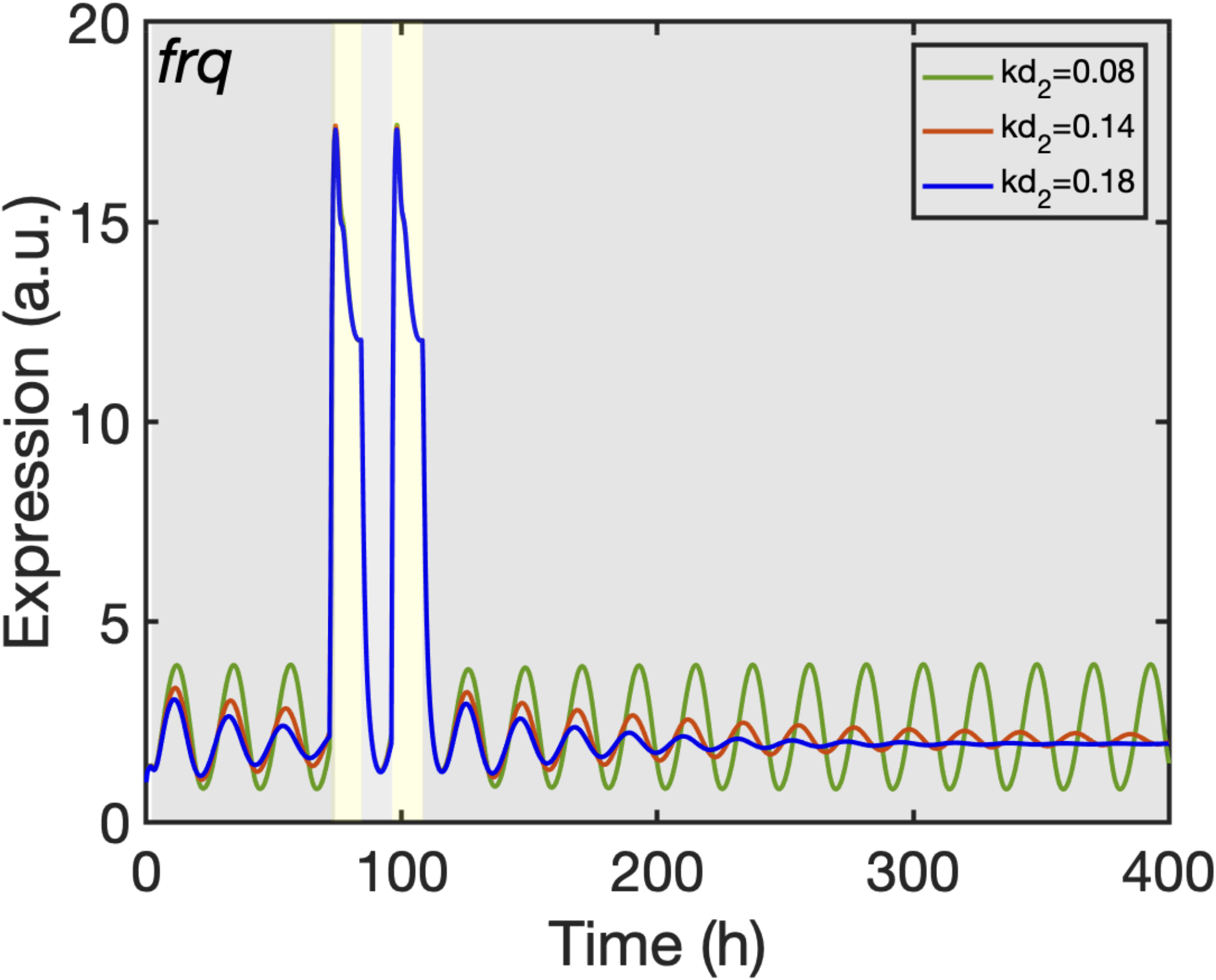
Impact of WCC dephosphorylation rate on frq expression rhythm. The modeled *frq* expression rhythm damps in the dark (blue trajectory). When the predicted dephosphorylation (reactivation) rate of WCC (k_d2_ = 0.18) was stepwise lowered dampening was reduced (k_d2_ = 0.14, red trajectory) or abolished (kd2 = 0.08, red trajectory) and the amplitude increased.

