## Supplementary file for "Data-driven modelling captures dynamics of the circadian clock of *Neurospora crassa*"

### *Supplementary Material*: Data-driven modelling of circadian clock of *Neurospora crassa*

#### 1 MATHEMATICAL MODEL

##### WCC dynamics

The white collar-1 (WC-1) and white collar-2 (WC-2) protein forms a white collar complex (WCC) (Talora et al., 1999). WCC senses the light through the N-terminal of PAS domain and LOV domain of WC-1 that subsequently changing the activity of WCC from dark to light, and form WCC homodimer. (Ballario et al., 1998; Froehlich et al., 2002; Cheng et al., 2003; Chen et al., 2010; Wu et al., 2014). Due to the presence of flavin adenine dinucleotide (FAD)-binding photoreceptor in WC-1, the activation of WCC differ from dark to light condition (He et al., 2002; Cheng et al., 2003; Wang et al., 2014). The VVD protein which contains LOV domain behave as a both positive regulator as well as inhibitor on WCC and form WCC-VVD heterodimer complex in the presence of light (Gin et al., 2013). Here, we modelled *wc-1* mRNA (assuming WC-2 is sufficiently abundant) and four distinct activation forms of WCC protein, i.e. WCC expression in constant dark $W_d$, monomer form $W_l$, dimer form $W_lW_l$ and heterodimer complex form with VVD $W_lV_l$ in light.

##### Dynamics of dark form WCC

We translate the following aspects of dark-form WCC dynamics into three ordinary differential equations. The detailed descriptions for the model variables and parameters are given in Table S1 and S2.

1. Basal level of mWCC activity in dark, $W_lW_l$ mediated transcription followed by mCSP1 inhibition
2. Translation of $W_d$ protein and degradation
3. Inhibition of $W_d$ by FFC-mediated phosphorylation generating the negative feedback loop as well as dephosphorylation of $W_d$
4. Degradation of $W_d$ and $W_{dp}$

$$\frac{d[mWCC]}{dt} = K_{\text{basal}} + k_1 \frac{[W_lW_l]}{(K_1 + [W_lW_l])} \frac{K_{\text{cp}_1}}{(K_{\text{cp}_1} + [CSP])} - k_{\text{d1}}[mWCC] \tag{S1}$$

$$\begin{aligned} \frac{d[W_d]}{dt} &= k_2[mWCC] - k_{\text{pos1}} \frac{[W_d][FFC]}{(K_{\text{pos1}} + [W_d])} + k_{\text{dpos1}}[W_{dp}] - k_{\text{d2}}[W_d] \\ &\quad + k_4[W_lW_l] + k_5[W_lV_l] - l_1[light][W_d] + k_3[W_l] \end{aligned} \tag{S2}$$

$$\begin{aligned} \frac{d[W_{dp}]}{dt} &= k_{\text{pos1}} \frac{[W_d][FFC]}{(K_{\text{pos1}} + [W_d])} - k_{\text{dpos1}}[W_{dp}] - k_{\text{dp1}}[W_{dp}] \\ &\quad - l_1[light][W_{dp}] + k_3[W_{lp}] + k_5[W_{lp}V_l] + k_4[W_lW_{lp}] \end{aligned} \tag{S3}$$

##### Dynamics of light form WCC

1. Activation of dark form WCC by light (converting to light form WCC).
2. Inhibition of $W_l$ by FFC-mediated phosphorylation generating the negative feedback loop and de phosphorylation of $W_{lp}$

3. Degradation of $W_l$ and $W_{lp}$
4. Photo adduct decay of $W_l$ and $W_{lp}$

$$\frac{d[W_l]}{dt} = l_1[light][W_d] - k_3[W_l] + k_7[W_lW_l] - k_6[W_l]^2 - k_{\text{pos2}}\frac{[W_l][FFC]}{(K_{\text{pos2}} + [W_l])} + k_{\text{dpos2}}[W_{lp}] - k_{\text{d3}}[W_l] + k_8[W_lV_l] - k_9[W_l][V_l] \tag{S5}$$

$$\frac{d[W_{lp}]}{dt} = k_{\text{pos2}}\frac{[W_l][FFC]}{(K_{\text{pos2}} + [W_l])} - k_{\text{dpos2}}[W_{lp}] - k_{\text{dp2}}[W_{lp}] + l_1[light][W_{dp}] - k_3[W_{lp}] + k_8[W_{lp}V_l] - k_9[W_{lp}][V_l] - k_6[W_{lp}]^2 + k_7[W_lW_{lp}] \tag{S6}$$

#### WCC homodimer dynamics

1. Reversible dimerization of WCC ($W_lW_l$) in the presence of light
2. Inhibition of $W_lW_l$ by FFC-mediated phosphorylation, generating the negative feedback loop and de phosphorylation of $W_lW_{lp}$
3. Photo adduct decay of $W_lW_l$ and $W_lW_{lp}$
4. Degradation of $W_lW_l$ and $W_lW_{lp}$

$$\frac{d[W_lW_l]}{dt} = k_6[W_l]^2 - k_7[W_lW_l] - k_4[W_lW_l] - k_{\text{pos3}}\frac{[W_lW_l][FFC]}{(K_{\text{pos3}} + [W_lW_l])} + k_{\text{dpos3}}[W_lW_{lp}] - k_{\text{d4}}[W_lW_l] \tag{S7}$$

$$\frac{d[W_lW_{lp}]}{dt} = k_{\text{pos3}}\frac{[W_lW_l][FFC]}{(K_{\text{pos3}} + [W_lW_l])} - k_{\text{dpos3}}[W_lW_{lp}] - k_{\text{dp3}}[W_lW_{lp}] + k_6[W_{lp}]^2 - k_7[W_lW_{lp}] - k_4[W_lW_{lp}] \tag{S8}$$

#### WCC-VVD heterodimer dynamics

1. WCC and VVD heterodimerize ($W_lV_l$) in the presence of light, forming a negative feedback loop.
2. Inhibition of $W_lV_l$ by FFC-mediated phosphorylation generating the negative feedback loop and de phosphorylation of $W_{lp}V_l$
3. Photo adduct decay of $W_lV_l$ and $W_{lp}V_l$ heterodimer complex
4. Degradation $W_lV_l$ and $W_{lp}V_l$

$$\frac{d[W_lV_l]}{dt} = k_9[W_l][V_l] - k_8[W_lV_l] - k_{\text{pos4}}\frac{[W_lV_l][FFC]}{(K_{\text{pos4}} + [W_lV_l])} + k_{\text{dpos4}}[W_{lp}V_l] - k_{\text{d5}}[W_lV_l] - k_5[W_lV_l] \tag{S9}$$

$$\frac{d[W_{lp}V_l]}{dt} = k_{\text{pos4}}\frac{[W_lV_l][FFC]}{(K_{\text{pos4}} + [W_lV_l])} - k_{\text{dpos4}}[W_{lp}V_l] - k_{\text{dp4}}[W_{lp}V_l] - k_8[W_{lp}V_l] + k_9[W_{lp}][V_l] - k_5[W_{lp}V_l] \tag{S10}$$

#### VVD dynamics

$W_l$ binds to the LRE region of the *vvd* gene and increases it's activity (Heintzen et al., 2001). The VVD protein disrupts the WCC homodimerization by competitive binding to WC-1 in the presence of light which acts as a feedback inhibitors of WCC (Chen et al., 2010; Hunt et al., 2010; Malzahn et al., 2010; Gin et al., 2013).

1. *vvd* transcription regulated by $W_d$, $W_l$, $W_lW_l$ and $W_lV_l$
2. Translation of *vvd* mRNA and VVD protein degradation
3. Conversion of $V_d$ to $V_l$ in the presence of light and degradation
4. Conversion of $V_l$ to $V_lV_l$ and degradation

$$\frac{d[mVVD]}{dt} = v_1\frac{[W_d]}{(K_2+[W_d])} + v_1\frac{[W_l]}{(K_2+[W_l])} + v_2\frac{[W_lW_l]}{(K_3+[W_lW_l])} + v_1\frac{[W_lV_l]}{(K_2+[W_lV_l])} - k_{\mathrm{d6}}[mVVD] \quad \text{(S11)}$$

$$\frac{d[V_d]}{dt} = v_3[mVVD] - l_1[light][V_d] + k_{10}[V_l] - k_{\mathrm{d7}}[V_d] \quad \text{(S12)}$$

$$\begin{aligned}\frac{d[V_l]}{dt} &= l_1[light][V_d] - k_{10}[V_l] + k_{11}[V_lV_l] - k_{12}[V_l]^2 - k_{\mathrm{d8}}[V_l] \\ &+ k_8[W_lV_l] - k_9[W_l][V_l] + k_5[W_lV_l] + k_5[W_{lp}V_l] + k_8[W_{lp}V_l] - k_9[W_{lp}][V_l]\end{aligned} \quad \text{(S13)}$$

$$\frac{d[V_lV_l]}{dt} = k_{12}[V_l]^2 - k_{11}[V_lV_l] - k_{\mathrm{d9}}[V_lV_l]$$

#### FRQ dynamics

The $W_d$ controls the transcription of the *frq* gene by binding to its clock box (C-box) in the dark which is essential and sufficient for rhythmic expression of *frq* (Froehlich et al., 2003). The WCC(L) binds to the light responsive element (LRE) of *frq* in the presence of light and promote the *frq* expression (Cheng et al., 2001; He and Liu, 2005; Dunlap, 2006). The FRQ protein phosphorylate and transport to the nucleus (Garceau et al., 1997; Luo et al., 1998; Liu et al., 2000; Diernfellner et al., 2009; Cha et al., 2011). WCC is hypophosphorylated and transcriptionally active in the absence of FRQ and shows opposite effect when FRQ is presence (Schafmeier et al., 2005). The FRQ protein inhibits WCC activity and show as a negative feedback loop (Aronson et al., 1994).

1. *frq* transcription regulated by $W_d$, $W_l$, $W_lW_l$ and $W_lV_l$
2. Translation of inactive form FRQ and its degradation
3. In general the conversion of inactive FRQ to FCC is a complex process which involve series of reactions. Here, we introduce five steps from inactive FRQ to mature FCC.
   $frqmRNA \xrightarrow{\text{v6}} FRQ \xrightarrow{\text{k13}} FRQ_1 \xrightarrow{\text{k13}} FRQ_2 \xrightarrow{\text{k13}} FRQ_3 \xrightarrow{\text{k13}} FRQ_4 \xrightarrow{\text{k13}} FRQ_5 \xrightarrow{\text{k13}} FFC \xrightarrow{\text{kd12}} \theta$ where the final FFC promotes the phosphorylation of WCC.

$$\frac{d[mFRQ]}{dt} = v_4\frac{[W_d]}{(K_4+[W_d])} + v_4\frac{[W_l]}{(K_4+[W_l])} + v_5\frac{[W_lW_l]}{(K_5+[W_lW_l])} + v_4\frac{[W_lV_l]}{(K_4+[W_lV_l])} - k_{\mathrm{d10}}[mFRQ] \quad \text{(S15)}$$

$$\frac{d[F_I]}{dt} = v_6[mFRQ] - k_{\mathrm{d11}}[FRQ] \quad \text{(S16)}$$

$$\frac{d[FRQ_1]}{dt} = k_{13}([FRQ] - [FRQ_1]) \quad \text{(S17)}$$

$$\frac{d[FRQ_i]}{dt} = k_{13}([FRQ_{i-1}] - [FRQ_i]), i = 2, ..., 5 \tag{S18}$$

$$\frac{d[FFC]}{dt} = k_{13}[FRQ_5] - k_{\mathrm{d12}}[FFC] \tag{S19}$$

##### 1.0.1 **CSP-1 dynamics**

CSP-1 is a global circadian repressor which is regulated by WCC, and it is essential for glucose compensation (Sancar et al., 2011). Additionally, CSP-1 inhibits the mRNA expression production of WCC that subsequently change the WCC activity in glucose dependent manner as well as it inhibits own gene expression (Sancar et al., 2011, 2012). Here, given the very short lifetime of csp-1 mRNA, we modelled the regulation of CSP-1 expression by $W_d$, $W_l$, $W_lW_l$ and $W_lV_l$ inhibition of its own production as following,

$$\begin{aligned} \frac{d[CSP]}{dt} &= v_7 \frac{[W_d]}{(K_6 + [W_d])} \frac{K_{\mathrm{cp2}}}{(K_{\mathrm{cp_2}} + [CSP])} + v_7 \frac{[W_l]}{(K_6 + [W_l])} \frac{K_{\mathrm{cp2}}}{(K_{\mathrm{cp_2}} + [CSP])} \\ &+ v_8 \frac{[W_lW_l]}{(K_7 + [W_lW_l])} \frac{K_{\mathrm{cp3}}}{(K_{\mathrm{cp_3}} + [CSP])} \\ &+ v_7 \frac{([W_lV_l]}{(K_6 + [W_lV_l])} \frac{K_{\mathrm{cp2}}}{(K_{\mathrm{cp_2}} + [CSP])} - k_{\mathrm{d13}}[CSP] \end{aligned} \tag{S20}$$

##### 1.0.2 **FAM-3 dynamics**

The CSP-1 effectively represses the transcription of *fam-3* (ncu09497) gene, so that *fam-3* expression shows anti-phasic behaviour compared to *frq* or *csp-1* (Sancar et al., 2011). Here, we assumed the basal level of transcriptional activation of *fam-3* is negatively regulated by followed by transcription inhibition by CSP-1.

$$\frac{d[mFAM]}{dt} = K_{\mathrm{basal1}} + v_9 \frac{K_{\mathrm{cp4}}}{(K_{\mathrm{cp_4}} + [CSP])} - k_{\mathrm{d14}}[mFAM] \tag{S21}$$

##### **Parameter estimation**

The main aims of parameter estimation is to find the optimal sets of parameter for a given set of measurement data points assuming the model M can describe the data sufficiently. To derive the unknown model parameters for the circadian clock, we used the standard method of minimizing the $\chi^2$ value (weighted sum of squared residuals),

$$\chi^2(\theta) = \sum_{i=1}^{N} \Big( \frac{y_i - y(t_i, \theta)}{\sigma_i} \Big)^2 \tag{S22}$$

where $\theta$ is the parameter vector; $y_i$ and $\sigma_i$ are measured mean value and standard deviation at time $t_i$; $y(t_i, \theta)$ is the simulated result by the model. The parameter inference was performed using the package Data2Dynamics (d2d)(Raue et al., 2015). We used the Latin Hyper Cube sampling (500 samplings) to scan the initial parameter space. Luciferase reporters data in WT and $\Delta vvd$ were used for estimation of the parameters (Table-S3, Table-S4).

Table S1: Model Variable

| No. | Variable description | Variables |
|---|---|---|
| 1 | WCC protein in dark | $W_d$ |
| 2 | WCC protein phosphorylation in dark | $W_{dp}$ |
| 3 | WCC$^\star$ monomer protein in light | $W_l$ |
| 4 | WCC$^\star$ monomer protein phosphorylation in light | $W_{lp}$ |
| 5 | WCC$^\star$-WCC$^\star$ homodimer protein | $W_lW_l$ |
| 6 | WCC$^\star$-WCC$^\star$ homodimer phosphorylation | $W_lW_{lp}$ |
| 7 | WCC$^\star$-VVD$^\star$ heterodimer | $W_lV_l$ |
| 8 | WCC$^\star$-VVD$^\star$ heterodimer phosphorylation | $W_{lp}V_l$ |
| 9 | VVD protein in dark | $V_d$ |
| 10 | Light-activated VVD protein | $V_l$ |
| 11 | VVD-VVD protein homodimer | $V_lV_l$ |
| 12 | FRQ protein in inactive form | FRQ |
| 13 | FRQ protein in active form | FFC |
| 14 | *mRNAwcc* | mWCC |
| 15 | *mRNAvvd* | mVVD |
| 16 | *mRNAfrq* | mFRQ |
| 17 | CSP-1 | CSP |
| 18 | *mRNAfam-3* | mFAM |

Table S2: Model parameters

| No. | Parameter | Description | Parameter value(a.u.) |
|---|---|---|---|
| 1 | $K_{\mathrm{b}asal}$ | Basal transcription rate of mWCC | 0.14 |
| 2 | $k_1$ | Max.rate of mWCC | 1.4 |
| 3 | $K_1$ | Half-maximal rate of mWCC transcription by $W_lW_l$ | 0.0024 |
| 4 | $K_{\mathrm{cp}_1}$ | mWCC transcription inhibited by mCSP-1 | 0.45 |
| 5 | $k_{\mathrm{d1}}$ | mWCC degradation | 2.5 |
| 6 | $k_2$ | $W_d$ translation | 5.2e+02 |
| 7 | $k_{\mathrm{pos1}}$ | Max.rate of $W_d$ phosphorylation | 8.7 |
| 8 | $K_{\mathrm{pos1}}$ | Half-maximal of $W_d$ phosphorylation | 7.6 |
| 9 | $k_{\mathrm{dpos1}}$ | Dephosphorylation of $W_{dp}$ | 0.19 |
| 10 | $k_3$ | Photoadduct decay of $W_l$ and $W_{lp}$ | 0.16 |
| 11 | $k_4$ | Photoadduct decay of $W_lW_l$ and $W_lW_{lp}$ | 0.38 |
| 12 | $k_5$ | Photoadduct decay of $W_lV_l$ and $W_{lp}V_l$ | 0.18 |
| 13 | $k_{\mathrm{d2}}$ | $W_d$ degradation | 0.18 |
| 14 | $k_{\mathrm{dp1}}$ | $W_{dp}$ degradation | 0.9 |
| 15 | $l_1$ | $W_l$ activated by light | 40 |
| 16 | $k_6$ | Homodimerization of $W_l$ and $W_{lp}$ | 12 |
| 17 | $k_7$ | Homodimerization dissociation of $W_lW_l$ and $W_lW_{lp}$ | 0.2 |
| 18 | $k_8$ | Heterodimerization dissociation of $W_lV_l$ and $W_{lp}V_l$ | 0.41 |

| | | | |
|---|---|---|---|
| 19 | $k_9$ | Heterodimerization formation of $W_lV_l$ and $W_{lp}V_l$ | 3.7 |
| 20 | $k_{\mathrm{d3}}$ | $W_l$ degradation | 1.5 |
| 21 | $k_{\mathrm{pos2}}$ | Max.rate phosphorylation of $W_l$ | 4.7 |
| 22 | $K_{\mathrm{pos2}}$ | Half-maximal rate of phosphorylation of $W_l$ | 3.5 |
| 23 | $k_{\mathrm{dpos2}}$ | Dephosphorylation of $W_{lp}$ | 0.13 |
| 24 | $k_{\mathrm{dp2}}$ | $W_{lp}$ degradation | 0.048 |
| 25 | $k_{\mathrm{pos3}}$ | Max.rate phosphorylation of $W_lW_l$ | 1.3 |
| 26 | $K_{\mathrm{pos3}}$ | Half-maximal rate of phosphorylation of $W_lW_l$ | 5.7 |
| 27 | $k_{\mathrm{dpos3}}$ | De phosphorylation of $W_lW_{lp}$ | 0.063 |
| 28 | $k_{\mathrm{d4}}$ | Degradation of $W_lW_l$ | 0.54 |
| 29 | $k_{\mathrm{dp3}}$ | Degradation of $W_lW_{lp}$ | 0.063 |
| 30 | $k_{\mathrm{pos4}}$ | Max.rate phosphorylation of $W_lV_l$ | 0.027 |
| 31 | $K_{\mathrm{pos4}}$ | Half-maximal rate of phosphorylation of $W_lV_l$ | 2.7 |
| 32 | $k_{\mathrm{dpos4}}$ | Dephosphorylation of $W_lV_{lp}$ | 1.7 |
| 33 | $k_{\mathrm{d5}}$ | $W_lV_l$ degradation | 6.1 |
| 34 | $k_{\mathrm{dp4}}$ | $W_lV_{lp}$ degradation | 0.063 |
| 35 | $v_1$ | Max. transcription of mVVD by $W_d$, $W_l$, $W_lV_l$ | 3.6 |
| 36 | $K_2$ | Half-maximal transcription of mVVD by $W_d$, $W_l$, $W_lV_l$ | 28 |
| 37 | $v_2$ | Max.rate transcription of mVVD by $W_lW_l$ | 97 |
| 38 | $K_3$ | Half-maximal transcription of mVVD by $W_lW_l$ | 2.1 |
| 39 | $k_{\mathrm{d6}}$ | mVVD degradation | 4* |
| 40 | $v_3$ | $V_d$ translation | 64 |
| 41 | $k_{\mathrm{d7}}$ | $V_d$ degradation | 2* |
| 42 | $l_1$ | Activation of $V_l$ by light | 40 |
| 43 | $k_{\mathrm{d8}}$ | $V_l$ degradation | 2* |
| 44 | $k_{10}$ | Photoadduct decay of $V_l$ | 0.2 |
| 45 | $k_{11}$ | Dimer formation of $V_lV_l$ | 3.6 |
| 46 | $k_{12}$ | Photoadduct decay of $V_lV_l$ | 0.34 |
| 47 | $k_{\mathrm{d9}}$ | $V_lV_l$ degradation | 2* |
| 48 | $v_4$ | mFRQ transcription by $W_d$, $W_l$, $W_lV_l$ | 11 |
| 49 | $K_4$ | Half-maximal transcription of mFRQ by $W_d$, $W_l$, $W_lV_l$ | 9.1e+02 |
| 50 | $v_5$ | mFRQ transcription by $W_lW_l$ | 4.6 |
| 51 | $K_5$ | Half-maximal transcription of mFRQ by $W_lW_l$ | 0.22 |
| 52 | $k_{\mathrm{d10}}$ | mFRQ degradation | 4* |
| 53 | $v_6$ | FRQ translation | 15 |
| 54 | $k_{\mathrm{d11}}$ | FRQ degradation | 0.8 |
| 55 | $k_{13}$ | FFC conversion | 1.02 |
| 56 | $k_{\mathrm{d12}}$ | FFC degradation | 0.67 |
| 57 | $v_7$ | mCSP-1 transcription by $W_d$, $W_l$, $W_lV_l$ | 1.5e+02 |
| 58 | $K_6$ | Half max transcription of mCSP-1 by $W_d$, $W_l$, $W_lV_l$ | 7.6e+02 |
| 59 | $v_8$ | mCSP-1 transcription by $W_lW_l$ | 46 |
| 60 | $K_7$ | Half max transcription of mCSP-1 by $W_lW_l$ | 7.9 |
| 61 | $K_{\mathrm{cp_2}}$ | mCSP-1 own transcription by $W_d$, $W_l$, $W_lV_l$ | 0.8 |

| | | | |
|---|---|---|---|
| 62 | $K_{\mathrm{cp}_3}$ | mCSP-1 own inhibition by $W_l W_l$ | 1.3 |
| 63 | $k_{\mathrm{d13}}$ | mCSP-1 degradation | 1.2 |
| 64 | $K_{\mathrm{basal1}}$ | mFAM-3 transcription in basal level | 0.001 |
| 65 | $v_9$ | mFAM-3 transcription | 3.9e+02 |
| 66 | $K_{\mathrm{cp}_4}$ | mFAM-3 transcription inhibited by mCSP-1 | 0.01 |
| 67 | $k_{\mathrm{d14}}$ | mFAM-3 degradation | 0.68 |

* Parameters are fixed.
